# DrugTar Improves Druggability Prediction by Integrating Large Language Models and Gene Ontologies

**DOI:** 10.1101/2024.09.21.614218

**Authors:** Niloofar Borhani, Iman Izadi, Ali Motahharynia, Mahsa Sheikholeslami, Yousof Gheisari

**Affiliations:** Department of Electrical and Computer Engineering, Isfahan University of Technology, Isfahan, Iran; Regenerative Medicine Research Center, Isfahan University of Medical Sciences, Isfahan, Iran; Isfahan Neuroscience Research Center, Isfahan University of Medical Sciences, Isfahan, Iran; Department of Medicinal Chemistry, Isfahan University of Medical Science, Isfahan, Iran; Department of Genetics and Molecular Biology, Isfahan University of Medical Sciences, Isfahan, Iran

**Author notes:** Corresponding Authors: Iman Izadi, Ph.D. Department of Electrical and Computer Engineering, Isfahan University of Technology, Isfahan, 84156-83111, Iran. Yousof Gheisari MD, PhD. Regenerative Medicine Research Center, Isfahan University of Medical Sciences, Isfahan, 81746-73461, Iran.

**Keywords:** Deep learning, Druggability prediction, Gene ontology, Large Language Model, Protein embedding, Target discovery

## Abstract

Target discovery is crucial in drug development, especially for complex chronic diseases. Recent advances in high-throughput technologies and the explosion of biomedical data have highlighted the potential of computational druggability prediction methods. However, most current methods rely on sequence-based features with machine learning, which often face challenges related to hand-crafted features, reproducibility, and accessibility. Moreover, the potential of raw sequence and protein structure has not been fully investigated. Here, we leveraged both protein sequence and structure using deep learning techniques, revealing that protein sequence, especially pre-trained embeddings, is more informative than protein structure. Next, we developed *DrugTar*, a highl7lperformance deep learning algorithm integrating sequence embeddings from the ESM-2 pre-trained protein language model with protein ontologies to predict druggability. DrugTar achieved areas under the curve and precision-recall curve values above 0.90, outperforming state-of-the-art methods. In conclusion, DrugTar streamlines target discovery as a bottleneck in developing novel therapeutics.

## Introduction

Drug development for complex chronic diseases is a challenge of current medicine. It is a costly, time-consuming process with a high failure rate, often due to the selection of incorrect targets for therapeutic intervention^1^. Hence, the primary focus in drug development lies in target discovery and validation. Despite the development of numerous FDA-approved drugs, these drugs target a minority of human proteins. Although a single protein can be targeted by multiple drugs, nearly 90% of proteins are not targeted by any drugs^2,3^. This indicates that certain intrinsic characteristics, make some proteins more suitable as drug targets^4^. Predicting such a “druggability” score paves the way for identifying novel therapeutic targets. However, this task cannot be simply performed using a small set of protein features and relies on sophisticated methods that consider the overall effect of a variety of parameters^2^. The development of these computational algorithms for druggability prediction provides an alternative strategy for the traditional time- and resource-consuming experimental approach.

While numerous computational methods have been developed for drug repositioning^5,6^, few studies have focused on predicting novel drug targets. In our previous study^2^, we developed a machine learning method that utilized biochemical characteristics and network topology parameters to classify drug targets (DT) and non-drug targets (non-DT). Our findings indicated that biochemical characteristics are more informative for predicting druggability than network topology features. Most druggability prediction methods have been developed using machine learning techniques that rely on sequence-based features, like amino acid composition (AAC), as well as physicochemical properties such as hydrophobicity, polarity, and charge of amino acids extracted from sequences^7–10^. Ensemble learning methods, which combine multiple models using different machine learning techniques or protein features, have also been applied to improve druggability prediction^4,11,12^. For instance, the SPIDER method^12^ employs diverse sequence-based descriptors and well-known machine learning algorithms through stacked ensemble learning.

The advancement of natural language processing (NLP) and large language models (LLM) has enabled the learning of word embeddings, which enhances our understanding of biological sequence functions. However, applying NLP to protein sequences presents challenges, such as the lack of clear words and a wide range of sequence lengths^13^. To address these, Sun et al^14^ employed strategies such as single and 3-gram word2vec embeddings with different machine learning algorithms. Yu et al^15^ were the first to apply deep learning techniques, including convolutional neural networks (CNN) and recurrent neural networks (RNN), to predict druggability from protein sequences. Additionally, QuoteTarget method^16^ leveraged graph neural networks (GNN) and the pre-trained protein language model ESM1b to model protein sequences and predicted structures as graphs to enhance performance.

Despite the value of developed methods for predicting protein druggability, several significant barriers have limited their widespread adoption. These algorithms typically use sequence-based features, which do not fully leverage the complete sequence information, or rely on manual biological feature extraction, making the process labor-intensive and time-consuming. On the other hand, recent deep learning methods could hardly ever surpass the functionality of machine learning strategies for druggability prediction, potentially due to the small size of the datasets. Furthermore, the introduced techniques are often unavailable as online tools, limiting their usability in experimental pipelines and hindering the assessment of their performance by independent investigators. Given these challenges and the critical gap in drug development, there is a clear need for a more robust, high-performance, and accessible method for predicting the tendency of proteins to be targeted by future therapeutic molecules. In this study, using the advantage of integrating pre-trained protein sequence embedding and protein ontologies, we developed DrugTar, a deep learning algorithm that demonstrates impressive functionality, superior performance compared to state-of-the-art methods, and consistent robustness across various assessments. This method, available as an online tool, facilitates target discovery, particularly in drug development for complex disorders.

## Results

### Protein sequence is more informative than its 3D structure for druggability prediction

Both sequence and structural information can be exploited in the druggability prediction. Protein functions are directly dictated by their 3D structures^17^. Specifically, drugs bind to the 3D structure of their target proteins. Although protein sequence determines almost all protein characteristics, including the 3D structure, predicting structure from sequence requires sophisticated techniques. Therefore, we assumed it is more straightforward to directly utilize structural features in the druggability prediction rather than sequence data. Due to the lack of suitable datasets, the ProTar-I dataset was constructed, which includes refined structural information of 2,248 proteins. This dataset includes both the atomic coordinates of 3D structures and the sequence information. For each protein, the highest resolution protein data bank (PDB) file with the longest sequence length was obtained from the RCSB PDB database^18^, ensuring that the PDB file covers at least half of the full protein sequence. Proteins were isolated as monomers by excluding additional chains, water molecules, ions, ligands, and other heteroatoms. ProTar-I consists of PDB files for 1,124 DTs and 1,124 non-DTs. The DTs are FDA-approved drug targets, while the non-DTs are not targeted by any FDA-approved or experimental drugs. Specifications of ProTar-I are provided in Supplementary Table 1.

To convert protein geometric 3D structures into a suitable format for deep learning methods, PDB files can be processed through several ways. One effective method is using point clouds, which reduce data volume compared to voxel grids. A point cloud consists of spatial data points extracted from PDB files, representing the 3D coordinates of atoms or amino acids. Here, amino acids were considered as points, defined by their alpha carbon coordinates. Additionally, amino acid coordinates were centralized to standardize protein positions. PointNet, a deep neural network designed to process 3D point clouds directly^19^, was utilized with the constructed protein point clouds to forecast druggability, referred to as “PointNet-PC”. Since PointNet requires a fixed number of points, a random set of 400 amino acids was selected for each protein with more than 400 amino acids, providing the overall geometric shape of the protein. The number 400 was chosen because it is close to the average length of the ProTar-I samples. Furthermore, zero padding was used for proteins with fewer than 400 amino acids. To increase the training dataset threefold and ensure the model is invariant to rotation, data augmentation was conducted by applying random rotations during training.

In addition to point clouds, a protein structure can be represented by the distance map, a symmetric matrix illustrating distances between residues. By applying a distance threshold, this matrix is transformed into a binary protein contact map. This 2D representation effectively encodes the protein structure^20^. In this study, a threshold of 7 Å was used to obtain a contact map, and a two-dimensional CNN was applied to predict druggability, referred to as “CNN2D- CM”. PointNet-PC and CNN2D-CM methods were evaluated using a 10-fold cross-validation (CV), and the functionality of the algorithms was assessed using various performance indices. The PointNet-PC and CNN2D-CM methods moderately discriminated between DTs and non- DTs, showing comparable performance (Fig. 1a). Given that structure-based methods did not yield satisfactory results, we turned to sequence-based methods.

**Figure 1.**
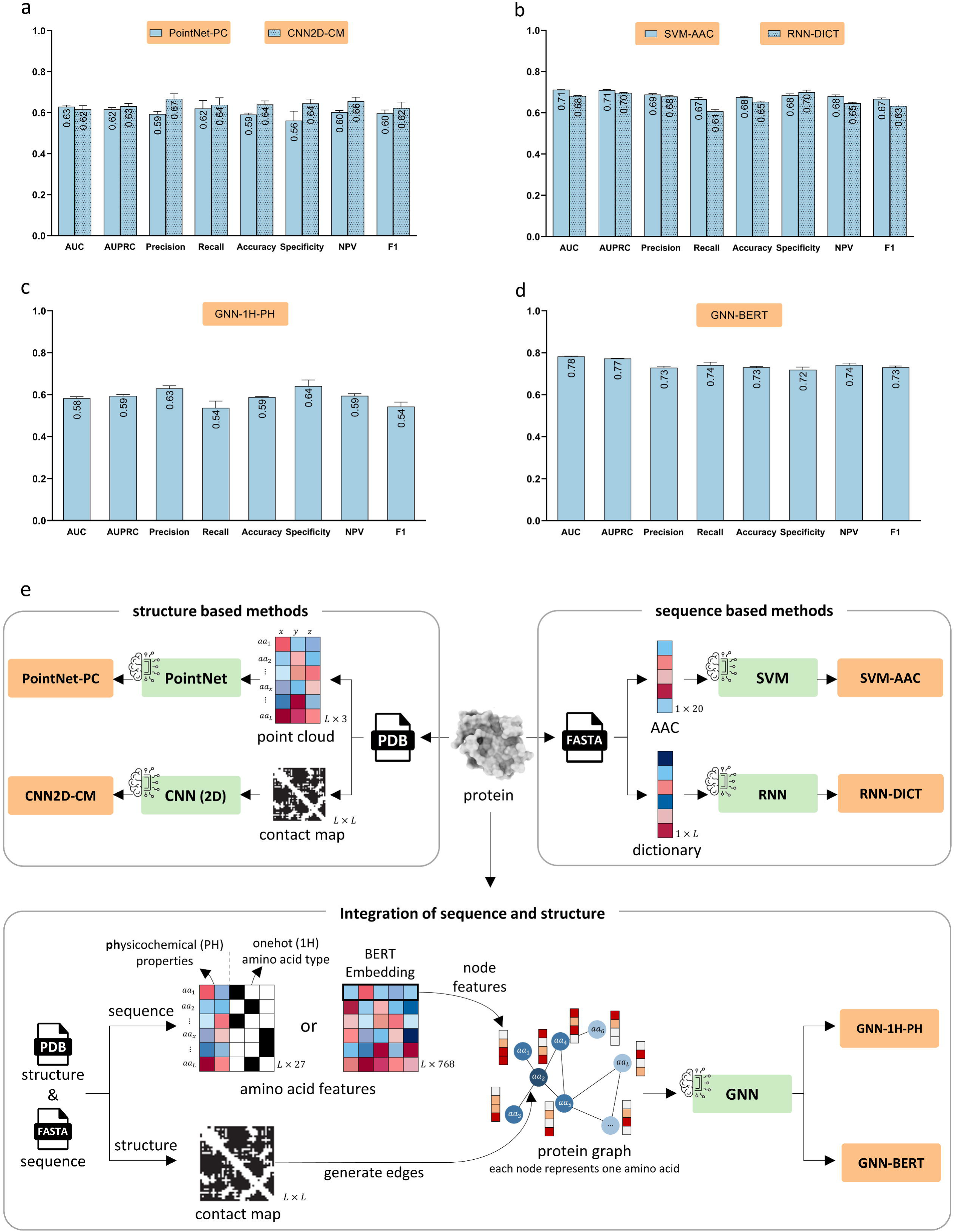
Comparison of sequence- and structure-based methods in druggability prediction. Both sequence- and structure-based approaches demonstrated moderate performance (a, b). The sequence and structure information were integrated with GNN models (c). Applying pre-trained BERT sequence embeddings as amino acid features led to improved performance in GNN-BERT compared to other methods, highlighting the role of sequence information (d). Schematics of implemented methods are illustrated in panel (e). Data are presented as the mean ± standard deviation from five independent runs, each using a 10-fold cross-validation method.

A variety of sequence features have been applied in the druggability prediction^10,12,21^, with one of the most common being amino acid compositions referred to as AAC. We began with AAC as a basic sequence feature and employed a support vector machine (SVM) as a simple classifier. This method is referred to as “SVM-AAC”. Although AAC is a common sequence feature, it is limited by its lack of amino acid order. To address this limitation, “RNN-DICT” was developed. This method incorporates sequence information through dictionary encoding and employs bidirectional long short-term memory (Bi-LSTM) with attention mechanisms. These sequence-based methods demonstrated moderate performance in druggability prediction (Fig. 1b). The AAC provides valuable information for predicting druggability despite its simplicity. Notably, these simple sequence-based algorithms slightly outperformed structure-based algorithms. It was observed that both sequence and structural information were moderately effective in discriminating between DTs and non-DTs.

To enhance performance, the next step was to integrate both sequence and structural data simultaneously. To this aim, a graph isomorphism network (GIN), a type of GNN known for its effectiveness in graph representation learning, was developed. For the first time, molecular graphs of proteins were constructed using real PDB files, with amino acids represented as nodes. The GIN propagates amino acid-level features across connected nodes based on the contact map matrix, generating protein-level feature representations for classification. Initially, one-hot encoding was used to represent amino acid types, along with seven physicochemical properties as outlined earlier^22^, specified as amino acid level features. The performance of this model referred to as “GNN-1H-PH”, was not satisfactory (Fig. 1c). Recent studies have shown that features obtained from pre-trained language models can significantly enhance classification performance^23^. Therefore, bidirectional encoder representations from transformers (BERT), a pre-trained language model specifically designed for protein sequences^24,25^, was utilized to obtain context-aware residue-level embedding vectors as node features. This method, referred to as “GNN-BERT”, leverages the power of pre-trained embeddings. While GNN-1H-PH showed insufficiently discriminative, GNN-BERT demonstrated significantly superior performance across different indices with the T-test P-value < 0.05 (Fig. 1d).

A Comparison between GNN-1H-PH and GNN-BERT models indicates that node features representing sequence information, significantly influence GNN performance. To assess the contribution of structural information, the GNN-BERT architecture and hyperparameters were kept consistent while retraining and assessing the model by replacing the contact map with a random undirected graph having an equivalent number of edges as the original, along with an identity matrix. Interestingly, the model performance did not decline when using the random graph or identity contact matrix compared to the original graph (Table 1). This finding suggests that structural information does not significantly contribute to the classification power of GNN- BERT, whereas the BERT sequence embedding features play a crucial role. Taken together, protein sequence is more informative than structure for druggability prediction

**Table 1.**
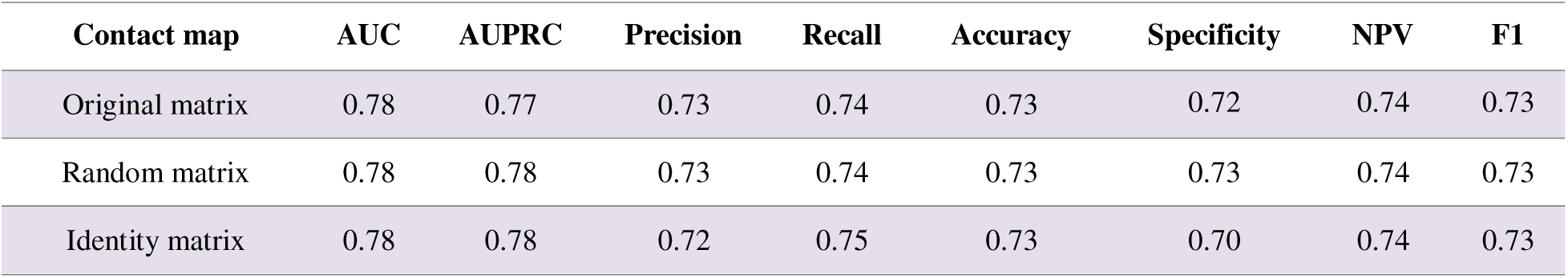
Outcomes for different types of structure by different contact maps in the GNN-BERT.

### Integrating pre-trained sequence embeddings with protein ontology data through deep learning enhances druggability prediction

To further assess the role of protein structure in druggability prediction, a method named DNN- BERT was proposed to compare with GNN-BERT. DNN-BERT operates by averaging the residue-level embedding vectors generated by BERT and utilizing them in a deep neural network for classifying DT and non-DT. Implementations on the ProTar-I dataset demonstrated that DNN-BERT, which relies solely on sequence information, performed similarly to GNN-BERT, which incorporates both sequence and structure data (Fig. 2a). This provides additional evidence that structural information does not offer insights beyond sequence embeddings. The advantage of DNN-BERT and similar models, which rely exclusively on sequence data is their broader applicability to proteins lacking structural data. This characteristic is valuable, as deep learning methods often yield better performance with larger datasets^26^. Hence, the ProTar-II dataset was constructed, containing sequence data for 2,034 DTs and 2,034 non-DTs, without any structural information (Supplementary Table 1). DNN-BERT showed strong performance on this dataset as well (Fig. 2a).

**Figure 2.**
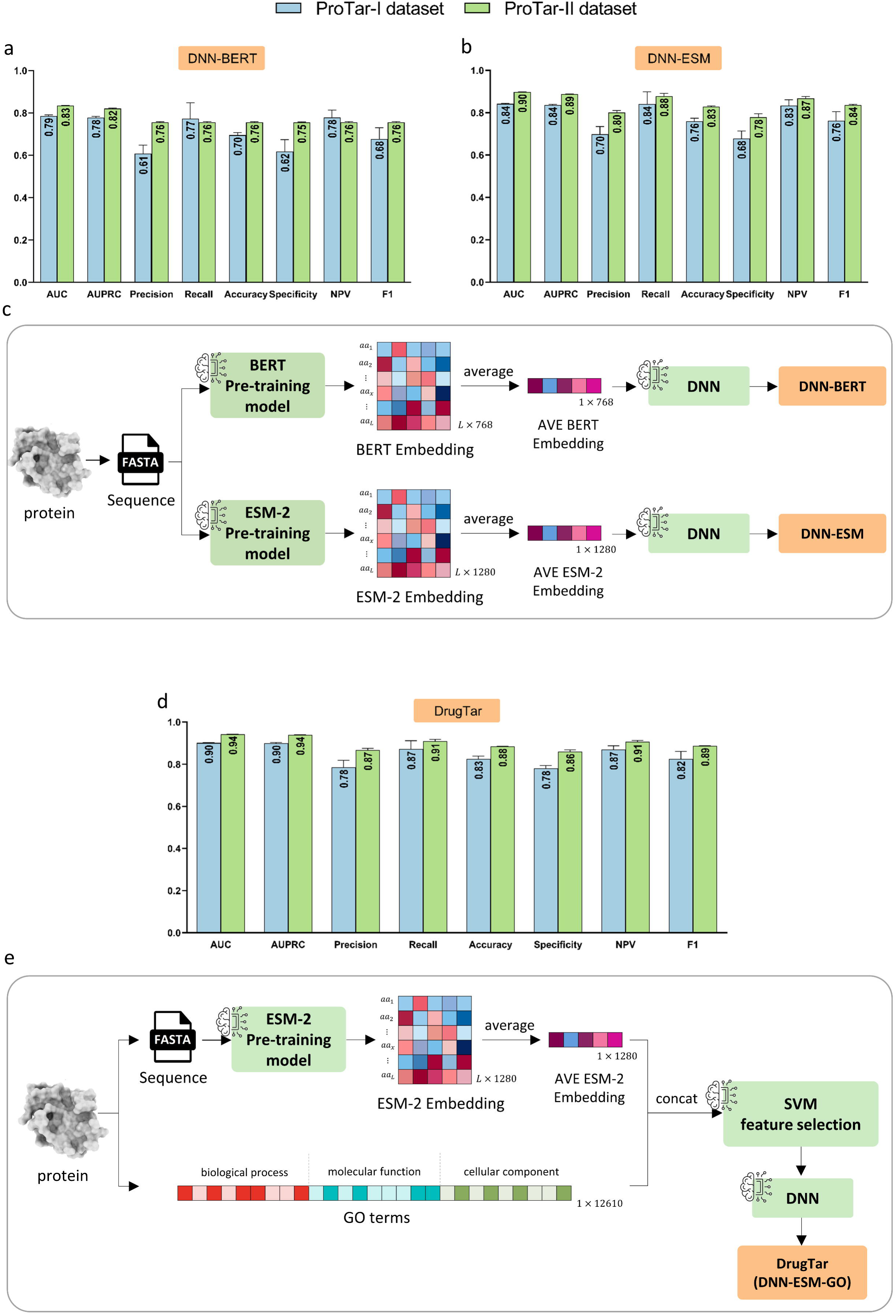
Enhancing druggability prediction with pre-trained sequence embeddings and gene ontologies. Using pre-trained sequence embeddings in a deep neural network yields satisfactory results. Despite only using sequence information, BERT-DNN achieved performance comparable to GNN-BERT, which requires both sequence and structure data, underscoring the importance of sequence information. The ESM-2 model surpassed BERT in druggability prediction on ProTar-I and ProTar-II datasets (a, b). The performance of the proposed DrugTar model for druggability prediction based on integrating pre-trained sequence embedding and gene ontologies showed significant improvements across both datasets (d). Method schematics are shown in panels (c) and (e). Data are presented as the mean ± standard deviation from five independent runs, each using a 10-fold cross-validation method.

As indicated, the superiority of the DNN-BERT method lies in its use of the BERT language model, which has 110 million parameters and is pre-trained on approximately 31 million protein sequences^24,25^. Hence, to further improve predictions, we focused on utilizing a more advanced sequence pre-training model. ESM-2 is a more advanced pre-trained method for embedding protein sequences than BERT, owing to its larger number of parameters and larger training dataset. The 33^rd^ layer of ESM-2, esm2_t33_650M_UR50D, with 650 million parameters, has been trained on approximately 65 million sequences^27^. The “DNN-ESM” was constructed similarly to DNN-BERT, with ESM-2 replacing BERT. The 10-fold CV measurements showed that DNN-ESM outperformed DNN-BERT across all metrics (p-value < 0.05, Fig. 2b). Despite the performance improvement, ESM-2 has a limitation due to its high RAM requirements when computing representations for long proteins. To address this, tokens exceeding 1024 in long protein were removed similar to the previous study^16^.

Previous investigations have indicated that DTs often share common biological properties^28^. Gene ontology (GO) terms describe different types of protein characteristics including molecular function, cellular component, and biological process. Hence, we explored the hypothesis that incorporating protein ontologies in the algorithm can enhance model performance. A new method, “DNN-ESM-GO”, was developed to integrate GO terms with ESM-2 embeddings. This method, renamed “DrugTar”, leverages the combined power of the ESM-2 embedding method and GO annotations within a deep neural network framework. In biological systems with high-dimensional features, effective feature selection is critical to improving classification accuracy and preventing overfitting^7^. Therefore, the SVM feature selection algorithm was employed on the concatenation of ESM-2 embedding and GO terms in DrugTar. Subsequently, a deep neural network with three hidden layers was utilized to predict protein druggability. DrugTar demonstrated robust performance on both the ProTar-I and ProTar-II datasets (Fig. 2d). The functionality of different algorithms was compared in the ProTar-II dataset, demonstrating the superiority of DrugTar with a p-value < 0.05 for all indices (Fig. 3a and b).

**Figure 3.**
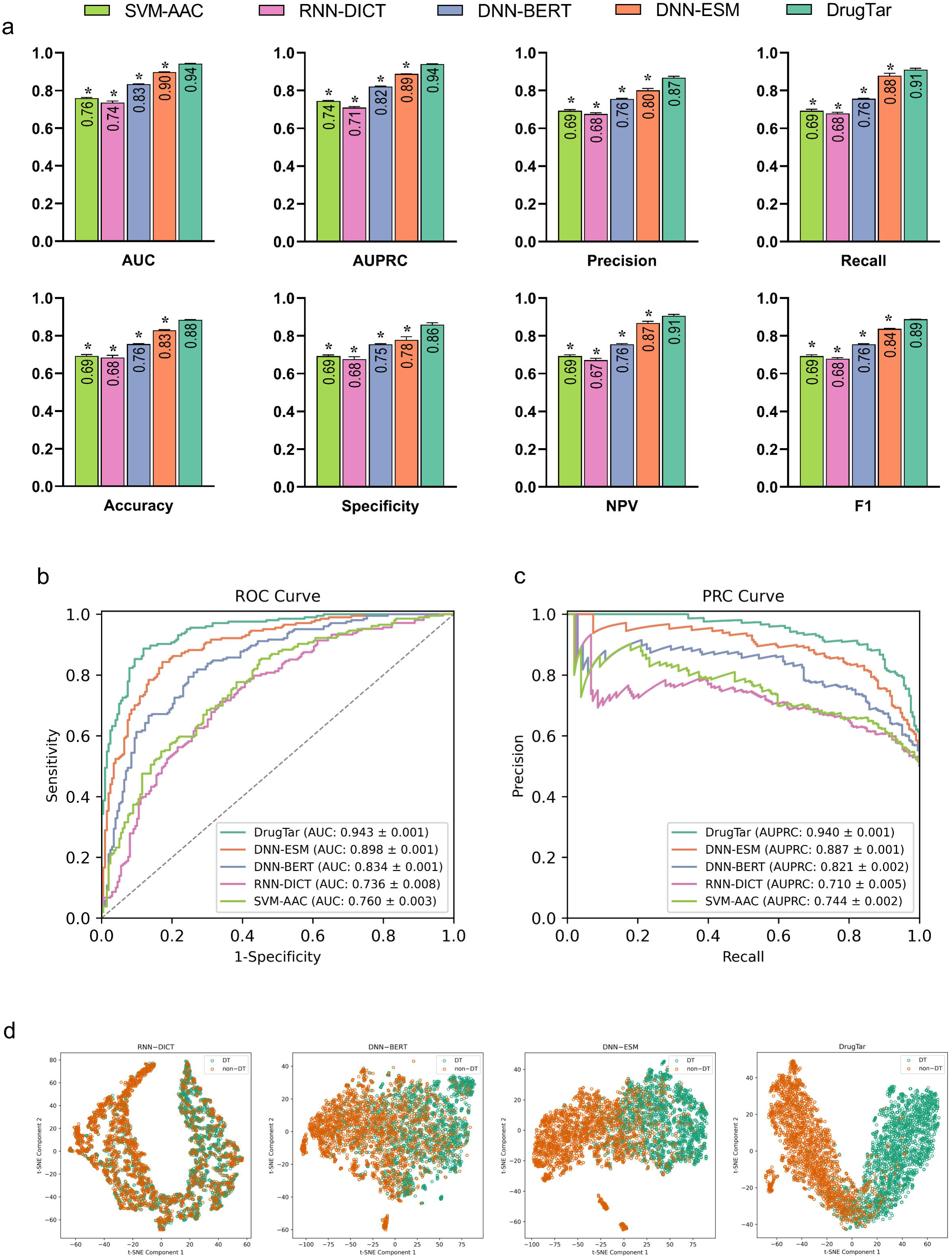
Performance evaluation of DrugTar and comparison with other methods on the ProTar-II dataset. The performance of the proposed DrugTar model was significantly superior to the examined models including sequence-based, GNN-1H-PH, GNN-BERT, DNN-BERT, and DNN-ESM (a). Performance is illustrated by the receiver operating characteristic (ROC) and precision-recall (PRC) curves, with each curve corresponding to a single run of 10-fold cross-validation (b, c). T-SNE visualization of protein latent vectors, where each marker represents a protein, showed that DrugTar provided superior discrimination between drug targets (DT) and non-drug (non-DT) targets proteins compared to other methods (d). Data are presented as the mean ± standard deviation from five independent runs.

To further evaluate the discriminative power of the methods, we applied the T-SNE (T-distributed stochastic neighbor embedding) algorithm^29^, a dimensionality reduction technique that maps similar objects in high-dimensional space into proximity within a reduced dimensional space. T-SNE was applied on the last dense layer before the output to visualize latent protein features in 2D space. The results showed that DrugTar provided better separation between DT and non-DT in the ProTar-II dataset compared to other methods, highlighting the superior predictive power of DrugTar (Fig. 3c).

### DrugTar demonstrates robustness and stability, outperforming state-of-the-art methods

To further evaluate the functionality and generalization of DrugTar, an independent test set, ProTar-II-Ind, consisting of 225 DT and 225 non-DT samples was created (Supplementary Table 1). DrugTar was initially trained and optimized using the ProTar-II dataset, after which it was applied to predict druggability scores for unseen samples in the ProTar-II-Ind. These scores were comparable to those obtained in the 10-fold CV on the ProTar-II dataset, demonstrating the strong generalization of DrugTar (Fig. 4a). Additionally, DrugTar outperformed other methods developed in the current study across all indices on the ProTar-II-Ind (P-value < 0.05).

**Figure 4.**
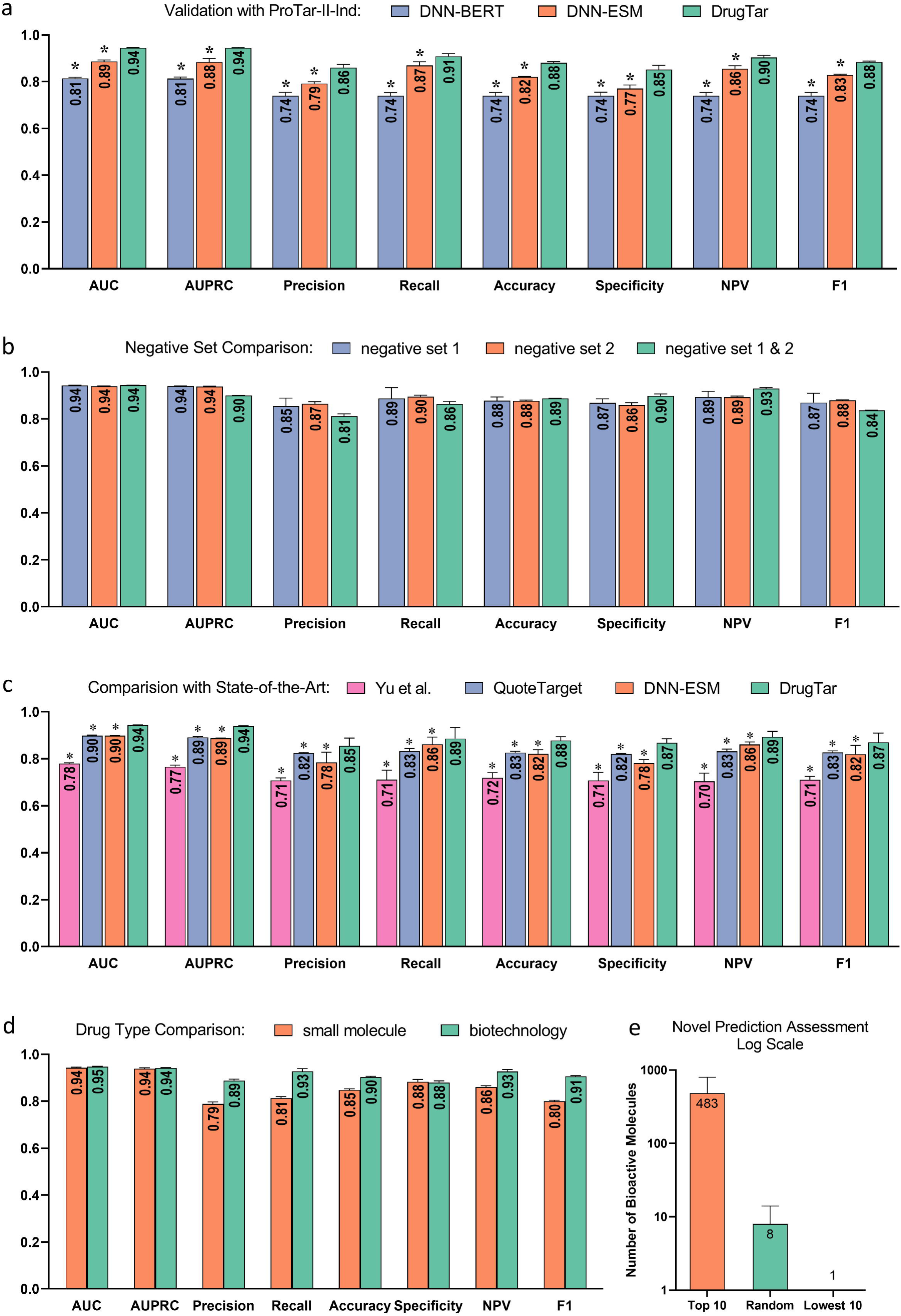
DrugTar demonstrated generalization and robustness in druggability prediction. DrugTar exhibited strong generalization and applicability on the independent test set ProTar-II- Ind (a). DrugTar showed stable and robust performance across various negative samples and imbalanced datasets (b). DrugTar consistently outperformed state-of-the-art methods on the ProTar-II dataset (c). DrugTar effectively predicts druggability scores for both biotechnology and small-molecule drug targets (d). The top 10 highest-scoring proteins, randomly selected sets of proteins, and the 10 lowest-scoring proteins predicted by DrugTar were validated using the ChEMBL database to assess the number of bioactive molecules, demonstrating the predictive power of the model (e). Data are presented as the mean ± standard deviation from five independent runs (a, b, c, d) and as the mean ± mean squared error (MSE) on a logarithmic scale (e).

The Jamali dataset, established in 2016, contains 1,224 DTs and 1,319 non-DTs^9^. It is widely used in druggability prediction studies^10–12,21^. The DTs in this dataset are FDA-approved drug targets, while the non-DTs are extracted using methods outlined by Li et al^30^ and Bakheet et al^31^, where relevant family members of DTs are excluded from the non-DT list. To analyze the Jamali dataset, the average AAC of the Jamali and ProTar-II datasets was compared, revealing significantly higher AAC differences between DTs and non-DTs in the Jamali dataset (Supplementary Fig. 1). This difference may be attributed to the method of selecting negative samples, suggesting that druggability prediction might be more straightforward in the Jamali dataset, potentially leading to overestimated performance. This underscores the significant impact of the method of selecting negative samples on the evaluation.

To account for this, a second negative set of 2,034 non-DTs, distinct from the negative set of ProTar-II, was extracted. DrugTar was evaluated using the same positive set but with this second negative set. The results showed comparable scores for both negative samples, indicating the robustness of DrugTar in handling negative samples. To further evaluate the robustness of DrugTar to an imbalanced dataset both negative sets were combined, creating an imbalanced dataset (2,034 DTs and 4,068 non-DTs). The DrugTar performance remained strong. To further evaluate the robustness of DrugTar to an imbalanced dataset, both negative sets were combined, creating an imbalanced dataset (2,034 DTs and 4,068 non-DTs). Despite the imbalance, the performance of DrugTar remained strong (Fig. 4b).

Even with the bias in negative sample selection in the Jamali dataset, DrugTar was evaluated against this dataset due to its frequent use by other methods. DrugTar demonstrated excellent performance, outperforming both SPIDER^12^ and Yu’s method^21^ (Table 2). However, considering the significant impact of the dataset used for training and evaluation in comparing deep learning methods, we proceeded to compare state-of-the-art methods using the ProTar-II dataset, ensuring a reliable and valuable comparison. Therefore, two current state-of-the-art methods were compared with DrugTar to verify that the features and network architecture employed by DrugTar provide superior predictions of protein druggability. QuoteTarget ^16^ and the best-performing method from the Yu et al study^32^, both of which are deep learning-based methods, were evaluated using a 10-fold CV on the ProTar-II dataset. DrugTar significantly outperformed these methods across all indices (P-value < 0.05, Fig. 4c). Interestingly, the performance of DNN-ESM was comparable to that of QuoteTarget, which feeds ESM1b sequence embedding and contact maps predicted by ESM1b into a GNN.

**Table 2.**
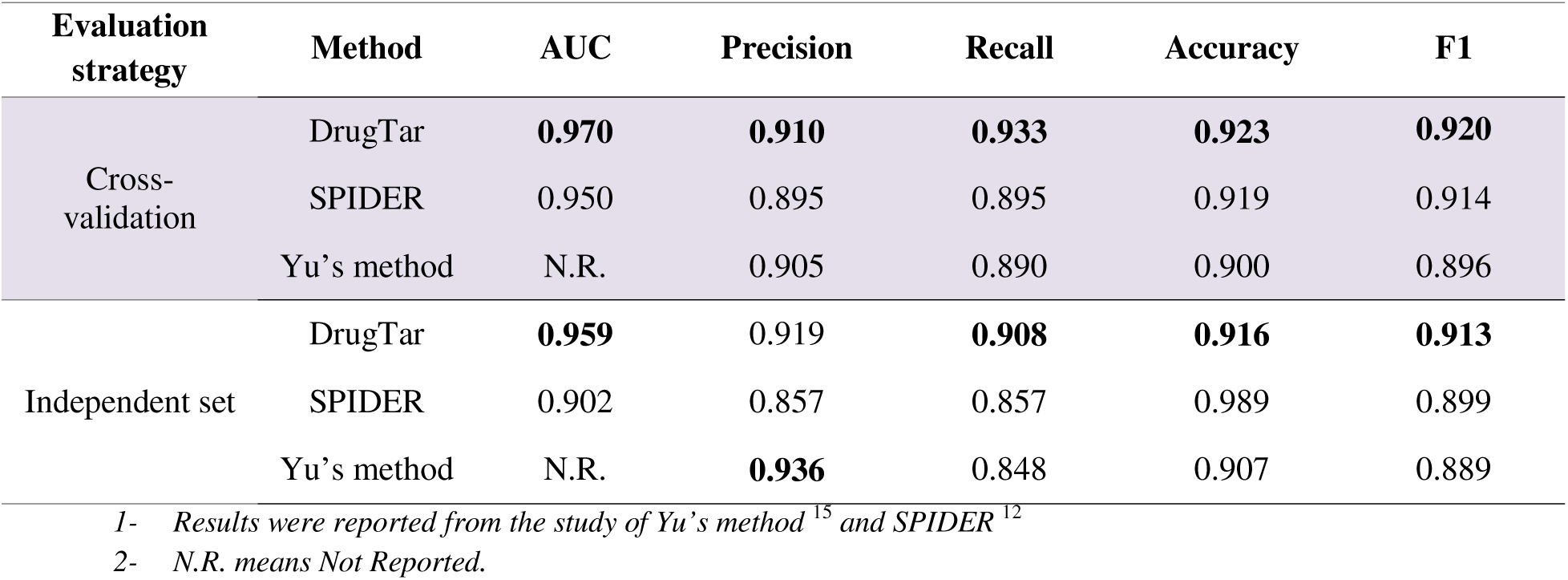
Performance comparison of DrugTar with state-of-the-art methods on Jamali Dataset.

DrugTar was also tested separately on targets for biotechnology and small-molecule drugs. The performance was satisfactory for both groups (P-value < 0.05, Fig. 4d), demonstrating that DrugTar effectively predicts druggability scores for both groups of drug targets. These results confirm that DrugTar is a robust and valid method for predicting druggability.

To assess the novel DrugTar predictions, we trained DrugTar on the ProTar-II dataset to predict druggability scores for proteins excluded from both the training and testing phases, ensuring all test proteins were unseen during model training and validation. We extracted the top 10 highest-scoring proteins, five randomly selected sets of 10 proteins, and the 10 lowest-scoring proteins as predicted by DrugTar. Subsequently, we checked the ChEMBL database^33^ to determine the number of bioactive molecules associated with these proteins. ChEMBL is a manually curated database of 2.4 million bioactive molecules with drug-like properties^34^. In this study, we consider the number of available bioactive molecules as an indicator of both the presence of an active site and the bioactivity of the target, with a higher count suggesting an increased likelihood of the target becoming a drug target. Among the top 10 highest-scoring proteins, 7 were found to be associated with several bioactive molecules. On average, these proteins have interactions with 482 ± 314 bioactive molecules. Notably, on average only 1 out of 10 proteins in the five random sets interacts with 8 ± 6 bioactive molecules and none of the 10 lowest-scoring proteins is associated with the bioactive molecules (Fig. 4e and Supplementary Tables 2, 3, and 4). These findings provide strong evidence that DrugTar effectively differentiates between highly druggable and less druggable targets, aligning well with empirical data and demonstrating its utility in prioritizing proteins for drug discovery.

## Discussion

Recent developments in high-throughput technologies have generated huge amounts of multi-omics data^35,36^. This has enabled the identification of extensive lists of proteins involved in various diseases, leading to the discovery of numerous proteins associated with disease pathways^37^. However, a significant challenge limiting the clinical translation of these findings is the difficulty in identifying which of the disease-associated genes are appropriate targets for therapeutic intervention. This study is dedicated to addressing this critical issue, aiming to bridge the gap between disease-associated gene discovery and clinical application. To achieve this, an innovative high-performance method named DrugTar was developed to predict the most druggable targets among a list of genes involved in disease pathogenesis.

To the best of our knowledge, this study is the first to compare sequence and structure information in druggability prediction using various algorithms. Remarkably, the results demonstrated that sequence data is more informative than the structure. This finding aligns with recent studies that reveal protein sequence, particularly sequence pre-trained embeddings, is more informative than protein structure for predicting protein function^38,39^, and protein interaction sites^40^. While structure-based methods are valuable, sequence-based models offer significant advantages, including enhanced accessibility and easier data processing. However, we appreciate that the superiority of sequence versus structure for druggability prediction remains limited to the employed methods and cannot be generalized.

The performance of SVM-AAC, as a simple machine learning method, was comparable to, and in some cases almost better than, sophisticated deep learning methods including GNN-1H-PH, and RNN-DICT. This is in line with the study of Yu et al who found that their deep learning method for druggability prediction was not better than machine learning algorithms^15^. This unexpected finding can be explained by the fact that machine learning often performs better on smaller datasets due to their simplicity and lower risk of overfitting. Conversely, as dataset size increases, deep learning models tend to excel because they can learn complex patterns from large amounts of data^41^. Labeled data is often scarce in biological settings, limiting the applicability of deep learning methods. Using pre-trained models on large, diverse datasets and then fine-tuning them on smaller, problem-specific datasets is a wise approach to this problem^23,42^. In this study, using pre-trained models like BERT or ESM-2 for sequence embedding significantly improved the distinction between DTs and non-DTs, consistent with previous findings that pre-trained sequence embeddings enhance performance in other biological applications^38,43^.

To enhance druggability prediction, DrugTar integrates ESM-2 sequence embeddings and protein ontologies through a deep neural network. Due to the large number of GO terms used in this algorithm, an SVM-based feature selection method was employed to avoid overfitting. Among the various available biological features, we chose to utilize protein ontologies as they are informative and represent intrinsic protein characteristics. Furthermore, they are disease-independent and can be easily and robustly accessed. Their hierarchical organization and well annotation provide additional advantages. The improvements achieved by incorporating GO terms are consistent with the findings of Balheet et al^31^ identifying that specific GO terms are associated with the tendency of proteins to be targeted by drugs. In line, we previously found that biochemical features of proteins are informative for druggability prediction^2^.

The appropriate selection of informative features in DrugTar, such as GO terms, allowed us to employ a simple classifier. Indeed, although we experimented with various sophisticated integration and classification algorithms, the simple 3-layer DNN implemented in DrugTar consistently delivered superior performance. This underscores the fact that simple classifiers with appropriate features can achieve excellent results^44^. Additionally, the simplicity of the classifier significantly reduces the computation cost.

The availability of a representative labeled dataset is a bottleneck in developing deep learning algorithms, particularly for classification. Any bias in the dataset may result in over- or underestimating the performance of the classifier. We have shown that a widely employed dataset in developing druggability prediction methods^9^ suffers from bias when selecting negative samples. Hence, the performance of the validated algorithms using this dataset remains to be further assessed with other datasets. In the present study, the functionality of DrugTar was assessed using different generated datasets including ProTar-I, ProTar-II, and ProTar-II-Ind. DrugTar consistently demonstrated a high performance using either of these datasets and surpassed state-of-the-art methods. Furthermore, the consistent performance across different negative sets demonstrates the robustness of DrugTar in handling negative samples. Additionally, when the size of negative samples was twice the size of positive samples, the performance of DrugTar remained high, indicating the robustness of this algorithm to unbalanced data. Additionally, DrugTar demonstrated reliable and discriminative performance both in the settings of small-molecule drugs and biotechnology pharmaceuticals targets. Notably, the validity of the top-ranked predictions was confirmed through empirical data from ChEMBL. These assessments clearly indicate the robust high performance of DrugTar for druggability prediction.

In conclusion, we developed DrugTar, a deep learning-based method that integrates ESM-2 sequence embedding with protein ontologies, achieving an impressive area under the ROC curve (AUC) and area under the precision-recall curve (AUPRC) of above 0.90. Extensive assessments across multiple datasets and comparisons with alternative tools demonstrated the robustness and effectiveness of DrugTar. The novel prediction capabilities of this method suggest its applicability in drug target prediction, particularly for complex disorders. By streamlining the selection of candidate proteins for pre-clinical examinations, DrugTar efficiently translates biological findings into clinical applications, thereby facilitating the drug development process. To enhance accessibility, we developed an online tool for DrugTar, available at http://DrugTar.com.

## Methods

### Dataset preparation

In this study, to assess various models, three datasets were created: the “ProTar-I” dataset, which includes PDB files representing protein structures; the “ProTar-II” dataset, consisting of FASTA files containing protein sequences; and the “ProTar-II-Ind” dataset, an independent set of FASTA files not included in ProTar-II, used to evaluate the generalization of the models trained with ProTar-II (Supplementary Table 1).

*ProTar-I.* To create the dataset capturing protein structure for drug target prediction algorithms, PDB files were obtained from the RCSB PDB database^18^, ensuring that the PDB file covers at least half of the full protein sequence. When multiple PDB files existed for a protein, the file with the longest residue length and the best resolution was selected (Supplementary Table 5). Since PDB files often contain multiple chains, we isolated monomer proteins by removing other chains. Furthermore, water molecules, ions, ligands, and other heteroatoms were eliminated. This dataset comprises 1,224 PDBs for DT proteins and 1,224 PDBs for non-DT proteins. The DTs are FDA-approved drug targets, while the non-DTs are not targeted by any FDA-approved or experimental drugs, according to the DrugBank database (version 5.1,^45,46^).

*ProTar-II and ProTar-II-Ind.* The approved DT human proteins were downloaded from the DrugBank database (version 5.1,^45,46^). Proteins not associated with drugs were obtained by excluding targets, enzymes, carriers, and transporters related to any drugs listed in the DrugBank database from a complete list of all reviewed human proteins (n = 20433) sourced from the UniProt database^47^. Out of 2,291 approved targets in DrugBank, 2,261 were found in the list of reviewed UniProt IDs, and their sequences were extracted. Then, proteins with sequence lengths smaller than 50 amino acids or greater than 10,000 amino acids were excluded. Employing negative sampling equal to 1 resulted in a final set comprising 2,259 DT proteins and 2,259 non-DT proteins. Protein sequences were extracted in FASTA format from the UniProt database. From this set, 90% of the data was used to construct ProTar-II, consisting of 2,034 DT proteins and 2,034 non-DT proteins (Supplementary Table 6). ProTar-II was split into 9:1 subsets for training and testing models using 10-fold cross-validation. The remaining 10%, consisting of 225 DTs and 225 non-DTs, was used to construct ProTar-II-Ind, a dataset for independent validation to assess model generalization (Supplementary Table 7). The GO annotation information, including biological process, molecular function, and cellular components was extracted from the GO database (gaf- version: 2.2,^48,49^). Annotations assigned to root terms, GO:0008150 (biological process), GO:0003674 (molecular function), and GO:0005575 (cellular component), were omitted to ensure the specificity.

### Structure- and sequence-based models

*PointNet-PC.* PointNet^19^ is an effective framework for processing raw point cloud data extracted from protein PDB files. Point order invariance and transformation invariance, are two essential properties of PointNet that make it highly compatible with the 3D structure of proteins. In this study, amino acids are considered as points, and their corresponding coordinates are the coordinates of the Cα atoms. We replicated the network architecture from the original paper^19^ and reduced the number of weights in each layer by half to better suit the druggability prediction and reduce overfitting (Supplementary Fig. 2).

*CNN2D-CM.* The CNN2D-CM model applies two-dimensional CNN on the protein residue contact map. To create contact maps, residues are considered to be in contact if the Euclidean distance between their corresponding Cα atoms is less than 10 Å. The CNN architecture consists of two convolutional layers, with filter sizes of 5×5×32 and 5×5×64, both using ReLU activation functions (Supplementary Fig. 3).

*SVM-AAC.* In the SVM-AAC model, the AAC vector, representing the frequency of 20 amino acids in the protein sequence, is used as an input feature for the SVM classifier to predict druggability.

*RNN-DICT.* RNNs are suitable for tasks where the order of information is crucial, such as sequence classification. Long short-term memory (LSTM) networks are particularly effective at retaining important information from earlier parts of the sequence, improving contextual understanding and predictions. The RNN-DICT method employs dictionary-encoded sequences and utilizes bi- LSTMs with attention mechanisms. A key feature of this model is an attention layer, which utilizes all output states from the bi-LSTMs to enhance prediction accuracy (Supplementary Fig. 4).

*Graph Isomorphism Network.* The GIN^50^ is an effective method for processing protein graphs and integrating both sequence and structural information. In this study, the molecular graph of proteins is constructed using real PDB files. To this aim, amino acids are defined as nodes and there is an edge between amino acids if the distance between their *C*_*α*_ atoms is less than 7 Å^51,52^. The GIN processes the adjacency matrix (contact map) along with amino acid features as input, learning a representation vector for both individual amino acids and the overall protein.

GIN iteratively updates the representation of each amino acid by aggregating the representations of its neighbors. Hence, the representation of the *i*_*th*_ amino acid obtained by the *k*_*th*_ layer of GIN is defined as follows:

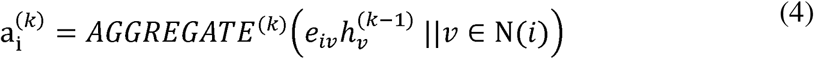

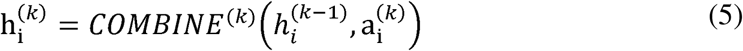

where Ν(*i*) denotes the neighbors of the *i*_*th*_ amino acid, and h is the representation of the *i*_*th*_ amino acid at the *k*_*th*_ GIN layer. Notably, the implemented method includes two GIN layers (*k* = 0,1,2), where ℎ^0^ refers to the initial amino acid feature. This feature can either be one-hot encoding with physicochemical properties in the GNN-1H-PH model, or BERT embeddings in the GNN- BERT model. The seven physicochemical properties of amino acids were outlined previously^22^. We opted for BERT embeddings over ESM-2 embeddings due to hardware limitations, as embedding all amino acids, especially in long proteins, would require significant RAM resources. The *e*_*iv*_ denotes the feature edge between the *i*_*th*_ and *i*_*th*_ amino acids, which are defined as:

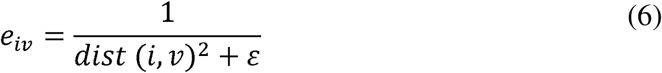

where *dist*(*i*, *i*) denotes the distance between the *C*_*α*_ atoms of the *i*_*th*_ and *i*_*th*_ residue, and ε is set to 10^−6^. For the *AGGREGATE*^(*k*)^ operation in Eq. 4, the sum operation is used, and for *COMBINE*^(*k*)^operation in Eq. 5, a linear function is applied, the same as in the previous study^50^. The overall protein representation *h*_*pr*_ is obtained by applying the mean function on the representations of all amino acids:

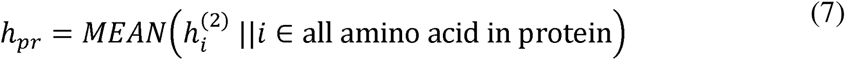

Where *h*^(2)^_*i*_ denotes the representation of the *i* amino acid at the second (final) GIN layer, and *i*_th_ represents the protein graph representation. Then, this vector is passed through two fully connected layers, with a final sigmoid activation function used to predict the druggability score (Supplementary Fig. 5).

*DNN-BERT and DNN-ESM.* The DNN-BERT method is based on a deep neural network that utilizes BERT embeddings of protein sequences. BERT^24,25^ is a pre-trained language model that generates context-aware residue-level embeddings, with each embedding vector having a size of 768 × 1. By averaging the embeddings of all amino acids in a protein sequence, the BERT protein representation is obtained. This representation is then input into a deep neural network for classifying DT and non-DT proteins (Supplementary Fig. 6). The DNN-ESM method follows a similar structure but utilizes ESM-2 for generating sequence embeddings instead of BERT. However, because BERT has lower memory requirements compared to ESM-2, all proteins in the dataset can be processed by the DNN-BERT method without the need to remove tokens.

### DrugTar structure

We developed a deep learning-based method called DrugTar, which integrated GO terms and ESM-2 embedding vectors through a deep neural network (Supplementary Fig. 7). DrugTar utilizes esm2_t33_650M_UR50D from ESM-2^27^, which was trained on nearly 650 million unique sequences, to extract amino acid representations. Protein sequences are fed into the pre-trained ESM-2 model, and the output from the 33^rd^ layer, consisting of 1,280 units, is extracted as the amino acid representation. For a protein of length *L*, the resulting sequence representation is a matrix *F* ∈ ℝ^1280×*L*^. Due to the high memory requirements for long protein sequences, tokens exceeding 1024 are removed. To compute the overall ESM-2 protein representation, the average of the amino acid embeddings is calculated as follows:

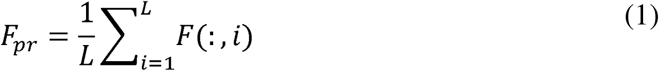

where *F*(: , *i*) represents the *i*_*th*_ amino acid representation, or the *i*_*th*_column of matrix *F*, and *F*_*pr*_ ∈ ℝ^1280×1^ is the resulting ESM-2 protein representation. To incorporate gene ontologies for druggability prediction, all three GO sub-ontologies, i.e., molecular function, cellular component, and biological process are encoded into a binary vector, denoted as *GO*_*pr*_ ∈ ℬ^12610×1^, and defined as follows:

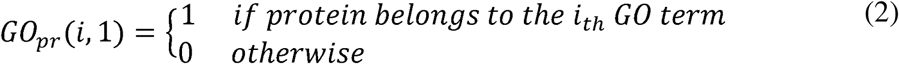

where *GO*_*pr*_ vector is encoding gene ontologies. After obtaining both the ESM-2 protein representation and GO binary vector, the protein representation is generated by concatenating them:

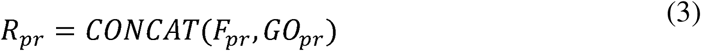

where *R*_*pr*_ is the protein representation used in the DrugTar algorithm. Performing feature selection is essential in biological systems characterized by high-dimensional features to enhance classification and prevent overfitting^7^. To address this, the SVM feature selection algorithm is applied, reducing *R*_*pr*_ dimension to 4000 × 1. This reduced representation is passed through a deep neural network consisting of three hidden layers. Finally, a sigmoid activation function is applied to the output neuron to predict a continuous druggability score for the protein, ranging from zero to one.

### DrugTar training and hyper-parameter tuning

The DrugTar method was implemented based on Tensorflow/Keras. It was trained by minimizing the average binary cross-entropy loss. The model was optimized using the Adam optimizer^53^. A custom learning rate scheduler was implemented, starting with an initial learning rate of 0.0002 for the first 5 epochs, which was halved every 5 epochs until reaching a minimum of 0.000025 after 15 epochs. The batch sizes were set to 32 for all datasets. The deep neural network architecture consisted of three hidden layers with 128, 64, and 32 units, respectively, using ReLU activation functions. Finally, a sigmoid function was applied in the final layer to predict a continuous druggability score between zero and one. To avoid overfitting, dropout^54^ with a drop probability of 0.5 was applied after the first and second dense layers, and L2 normalization with a penalty coefficient of 0.01 was used on the model weights. Furthermore, an early stopping criterion with the patience of 5 epochs was employed to optimize performance and prevent overfitting. The batch normalization was also applied after all dense layers.

### Model assessment

To assess the efficacy of the overall performance of various models, the 10-fold CV protocol was used. Each model was trained and tested 10 times, with each fold serving as the test set once, while the remaining nine folds were used for training. The ProTar-I and ProTar-II datasets were validated through this 10-fold CV process. For each fold, the performance of the models was assessed using eight metrics relevant to binary classification: AUC, AUPRC, precision, recall, accuracy, specificity, negative predictive value (NPV), and F1.

We also evaluate the generalization of models by constructing the ProTar-II-Ind dataset. The models were trained on the ProTar-II dataset and subsequently assessed using the independent test samples from the ProTar-II-Ind dataset. This approach was implemented to evaluate the generalization capability of the models, ensuring that the models perform well on unseen data.

### DrugTar web server

DrugTar is available as a web server at www.DrugTar.com. The web server enables users to predict druggability scores for proteins using the DrugTar method. Users can input protein IDs either by typing them directly or by uploading text or Excel files. The web server outputs DrugTar predictions and also offers downloadable documentation for users.

## Acknowledgment

Not applicable.

## Authors contribution

N.B., I.I., and Y.G. conceptualized the main idea. N.B., A.M., and M.S. gathered datasets. N.B. and I.I. proposed the method. N.B. performed simulations. N.B., I.I., and Y.G. contributed to the interpretation of the results. N.B., A.M., and M.S. drafted the manuscript and I.I., and Y.G. critically revised the manuscript. All authors approved the final draft and agreed to be responsible for the integrity of the entire work.

## Competing Interests

## Online Tool Website

http://DrugTar.com

## Code Availability

The source code related to the training and evaluation of the DrugTar model is publicly available with a detailed guide at https://github.com/NBorhani/DrugTar.

