## Supplementary Figures for "DrugTar Improves Druggability Prediction by Integrating Large Language Models and Gene Ontologies"

*** Corresponding Authors:**

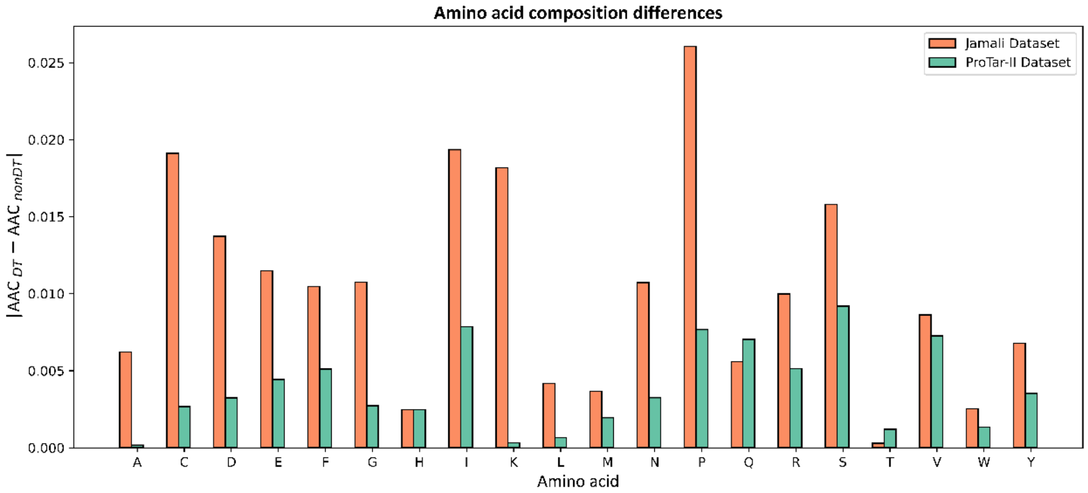

**Supplementary Figure 1- Differences in amino acid composition (AAC) between drug targets (DTs) and non-drug targets (non-DTs) in the Jamali and ProTar-II datasets.** The Jamali dataset exhibits significantly greater differences in AAC between DT and non-DT proteins compared to the ProTar-II dataset.

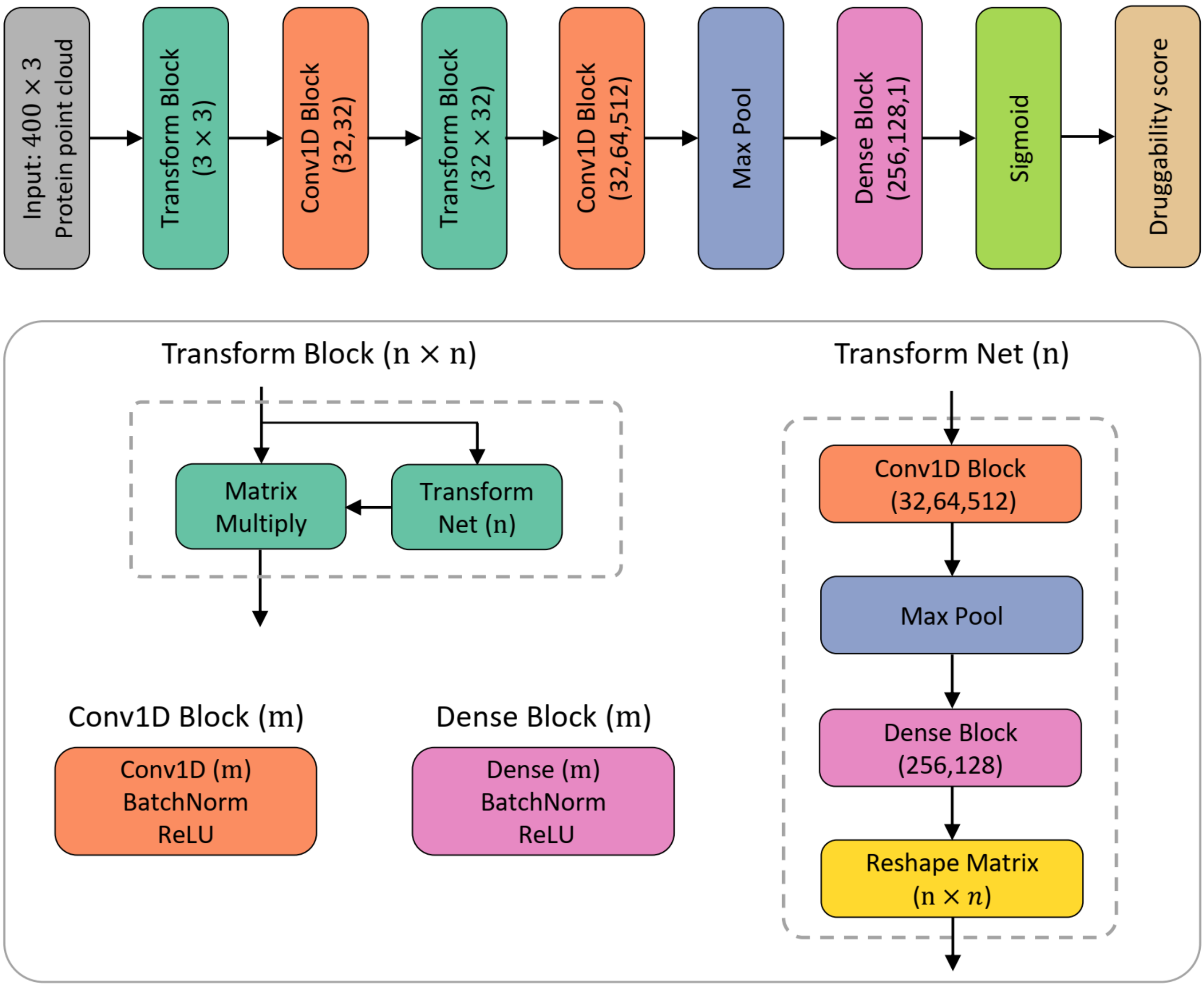

**Supplementary Figure 2- Overview of the PointNet-PC method framework.**

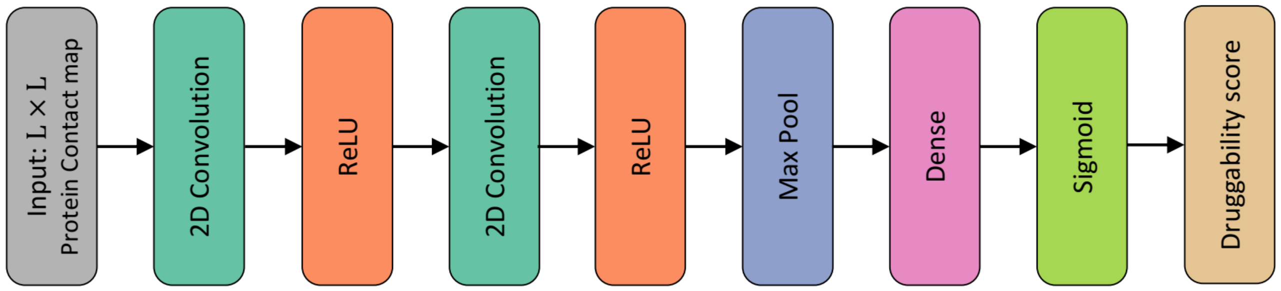

**Supplementary Figure 3- Overview of the CNN2D-CM method framework.**

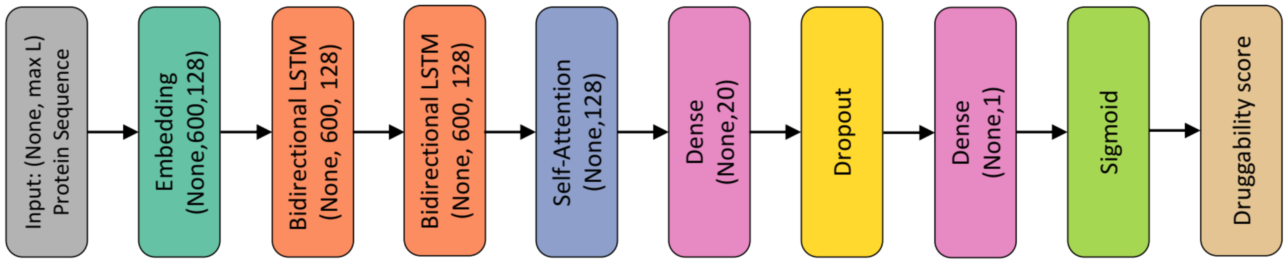

**Supplementary Figure 4- Overview of the RNN-DICT method framework.**

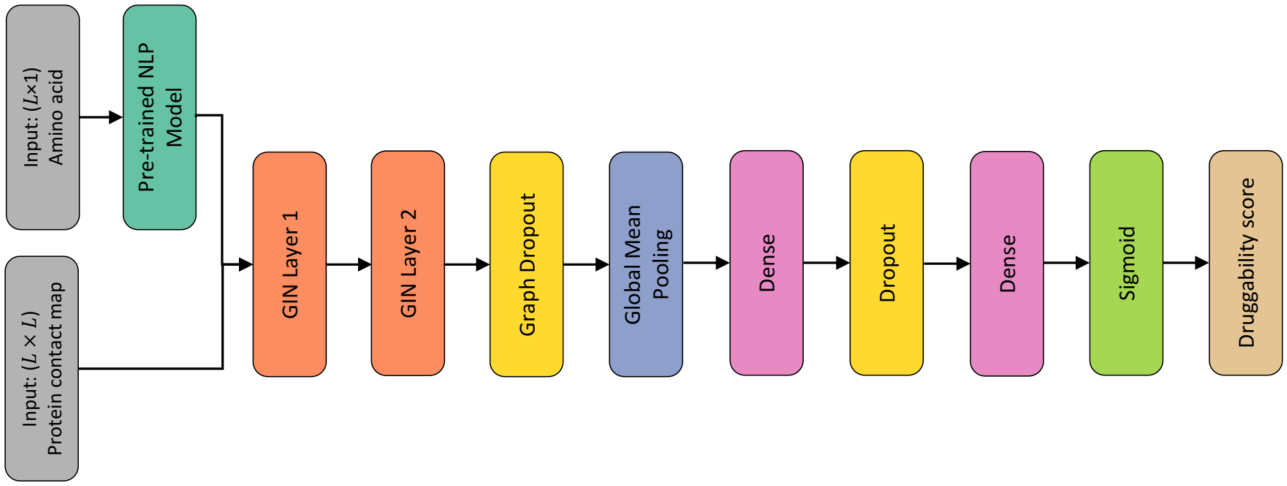

**Supplementary Figure 5- Overview of the GIN architecture in GNN-1H-PH and GNN-BERT method.**

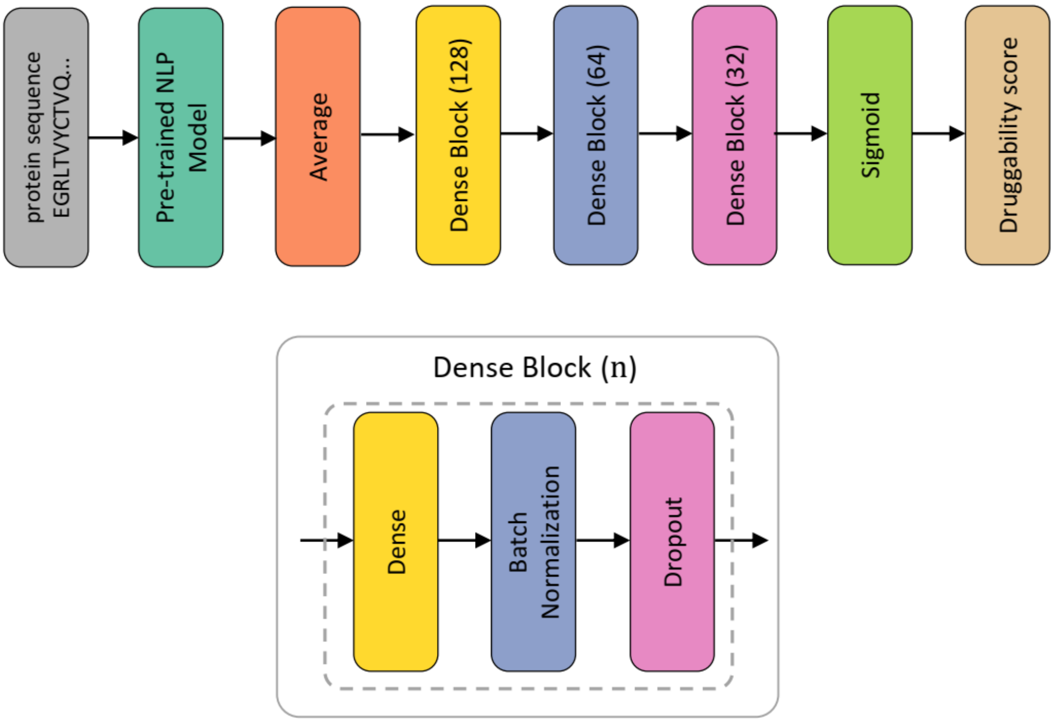

**Supplementary Figure 6- Overview of the DNN-BERT or DNN-ESM methods.**

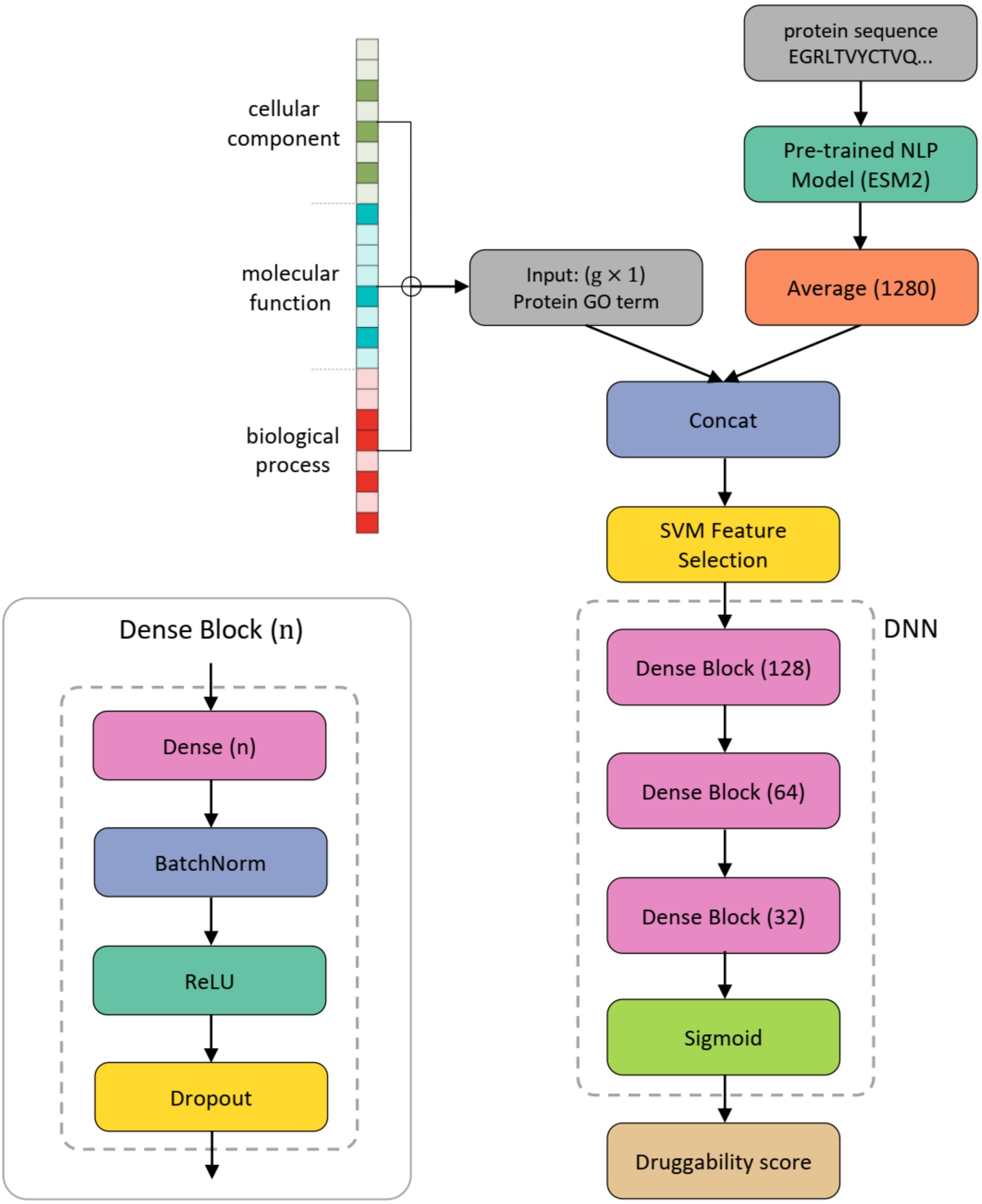

**Supplementary Figure 7- Schematic structure of DrugTar algorithm.**
